# Complement factor D (adipsin) mediates pressure-pain hypersensitivity post destabilization of medial meniscus injury

**DOI:** 10.1101/2025.05.29.656852

**Authors:** Priscilla M. Tjandra, Bethany A. Andoko, Jooyoung A. Kim, Andreana G. Gomez, Sonya Sar, Megha R. Aepala, Tiffany T.K. Pham, Darren Dumlao, Hope D. Welhaven, Kelsey H. Collins

## Abstract

**BACKGROUND:** Although osteoarthritis (OA) is the leading cause of pain and disability worldwide, there is a lack of models to probe the separable mechanism of OA structural damage and knee pain. We previously identified that deletion of complement factor D (FD) results in increased pressure-pain hyperalgesia despite cartilage protection after destabilization of the medial meniscus (DMM) surgery. However, how these discordant OA phenotypes manifest is not understood. We employed a novel targeted lipidomics approach to elucidate the role of eicosanoids in FD-mediated pain. We hypothesize that the absence of *Cfd (FD^−/−^)* will protect cartilage but cause increased pressure-pain hyperalgesia and eicosanoid dysregulation that persists throughout OA development.

**METHODS:** Male and female *FD^−/−^* and wild-type (WT) mice were challenged with DMM or remained naïve (n=5-11/group) at 16 weeks old. Pressure-pain hyperalgesia was measured bi-weekly for 8 weeks post-DMM. A second cohort was evaluated at 2 weeks post-DMM (n=6-10/group) to investigate DMM injury response. Structural damage was scored using the Modified Mankin system. To determine changes in eicosanoid profiles, serum and synovial fluid samples were analyzed via liquid chromatography-mass spectrometry (LC-MS). Statistical analysis was performed with unpaired t-test, two-way, or three-way ANOVA with Sidak’s posthoc test. Statistical significance is defined as p<0.05.

**RESULTS:** In contrast to WT mice, *FD^−/−^* showed no significant differences in Modified Mankin scores 8 weeks post-DMM. As expected, *FD^−/−^* hyperalgesia levels persisted until 8 weeks post DMM, similar to WT. Changes in eicosanoid profiles of pain-associated factors in *FD^−/−^*when compared to WT were found in the synovial fluid at 2 weeks and the serum at 8 weeks post-DMM.

**CONCLUSION:** The absence of *Cfd* drives knee hyperalgesia in male and female mice at 2 weeks-post DMM and persists through an 8-week observation period despite observing cartilage protection. Changes of eicosanoid profiles at both time points suggest that FD drives pain acutely, and the hyperalgesia phenotype emerges early in response to DMM injury, elucidating the role of the alternative complement in mediating OA pain and structural damage.

## INTRODUCTION

Osteoarthritis (OA), a disease characterized by the loss of cartilage lining the joint, is the leading cause of pain and disability worldwide^1,2^. Although pain is the primary driver for patients to seek care, current pain management strategies are inadequate, highlighting the need for new drug targets. However, disentangling the mechanism of OA pain from injury and disease entrenchment is challenging. For example, pain severity is not always a result of structural damage to the joint^3^. Until recently, most studies that employ preclinical models to study OA pathogenesis rely on structural characterization of the joint and often omit pain and behavioral changes. To address this gap in knowledge, we and others^4^ routinely perform pain phenotyping in all OA preclinical models to understand the concordant or divergent pain and structure phenotypes that manifest with OA in mice.

Complement factor D (FD), also known as adipsin, is a serine protease that cleaves complement factor B to activate alternative complement signaling, a key pathway in the innate immune response^5,6^. FD is primarily secreted from adipose tissue, which our lab has demonstrated is a key driver of OA pathogenesis and pain^7^. Although prior studies have determined that the loss of several complement signaling factors was protective for cartilage^8–10^, we have recently demonstrated that *FD^−/−^* mice displayed pronounced pressure-pain hyperalgesia, despite cartilage protection with DMM^11^. This model provides the opportunity to decode the discordance between structure and pain reported clinically and establishes FD as a key factor regulating fat-cartilage crosstalk in OA pain. In the present study, we leveraged this model of discordant pain and structural damage phenotype to investigate and better understand the time course changes and mechanism of pressure-pain hyperalgesia post-DMM.

Clinical data indicate that there is a dimorphism in OA, such that female patients can report more severe pain for a given amount of structural damage^12^. However, many in the field of OA believe female mice are a poor model to study OA, due to more modest structural changes in cartilage damage requiring a higher sample size to observe significant differences in histological OA scoring. Of note, several studies demonstrate that indeed female mice do develop hallmarks of OA in response to DMM^13,14^. Previously, increased synovial fluid C5 levels were associated with increased complement activation in male patients in late-stage knee OA compared to female patients^15^. Moreover, early complement activation has been reported to be higher in male humans and macaques^16^. These findings are also corroborated in mice and are thought to be due to restricted pathway components to promote inflammation through complement C5b-9 complex in female mice^17^. Therefore, in this study, we leverage the *FD^−/−^* model to gain insight into dimorphisms in pain regulation due to DMM, as it is known that innate differences in the modulation of pain – male mice may not adequately represent pain in female mice^4,18,19^. Therefore, a secondary aim of this study is to leverage the discordance in structural damage and pain in *FD^−/−^* mice to better understand potential sexually dimorphic pain responses due to DMM.

Current treatments for pain in early OA have been limited to non-steroidal anti-inflammatory drugs (NSAID)^20–22^. However, the efficacy of these drugs can be ineffective in OA patients for long-term use^23^. NSAIDs have been established as modulators of the complement system^24^. This same targeted eicosanoid panel^25^ demonstrated that alterations in the levels of several eicosanoid species played a key role in the transition from acute to chronic hypersensitivity in the K/BxN serum transfer model of rheumatoid arthritis through toll-like receptor (TLR) 4, another key regulator of innate immunity^26^. Previously, crosstalk between TLRs and complement signaling has been demonstrated to coordinate synergistic or exaggerated immune responses^27^. Studies have also shown that eicosanoids can be deployed downstream of complement signaling to assist with clearance and the inflammatory response to pathogens^28^.

To begin to elucidate the complement-mediated mechanisms that regulate pain in DMM-induced OA immediately after injury, we used a novel targeted approach in assessing eicosanoids, lipid-derived metabolites associated with pain, that are modulated by NSAIDs^25,29–31^ and can be downstream of complement signaling. As systemic soluble mediators appear to drive fat-cartilage crosstalk and OA after DMM^7^, we posit that *FD^−/−^*mice may demonstrate alterations in pain-driving and pain-relieving through eicosanoid mediators that act through these signaling molecules to modulate pain. Leveraging this novel eicosanoid profiling approach will give mechanistic insights to systemic and local cellular mechanisms related to pain that could lead to the development of novel therapeutic targets.

The purpose of this study was to determine the role of FD in the onset and persistence of pain in male and female mice acutely 2 weeks after DMM, and chronically after 8 weeks of DMM. We hypothesize that the absence of *FD^−/−^* will protect cartilage but result in pressure-pain hyperalgesia, which will be detectable 2 weeks post-DMM. We also hypothesize that hyperalgesia due to loss of FD would be more pronounced in male mice compared to female mice. As an exploratory hypothesis, we sought to test if eicosanoid dysregulation due to loss of FD is related to the pressure-pain hyperalgesia phenotype, potentially indicating novel fat- and lipid-derived targets that can be probed for the development of novel therapeutic strategies.

## METHODS

### Animal Studies

*FD^−^*^/−^ mice^6^ (provided kindly by J. Atkinson and X. Wu; Washington University in St. Louis) were bred and maintained at the animal facility at the University of California, San Francisco. All experimental procedures were approved by the University of California, San Francisco Institutional Animal Care and Use Committee. Male and female *FD^−/−^* and wild-type (WT) (Jackson Labs #005304) control mice were challenged with destabilization of the medial meniscus (DMM) on the left knee joints or remained naïve at 16 weeks old (n=5-11/group) as a control. Mice were sacrificed at either 2 weeks or 8 weeks post-DMM, 18 weeks or 24 weeks old in total age, at which time serum, synovial fluid, knee joints, and DRGs were collected. A timeline of these studies is presented in Figure 1a.

**Figure 1.**
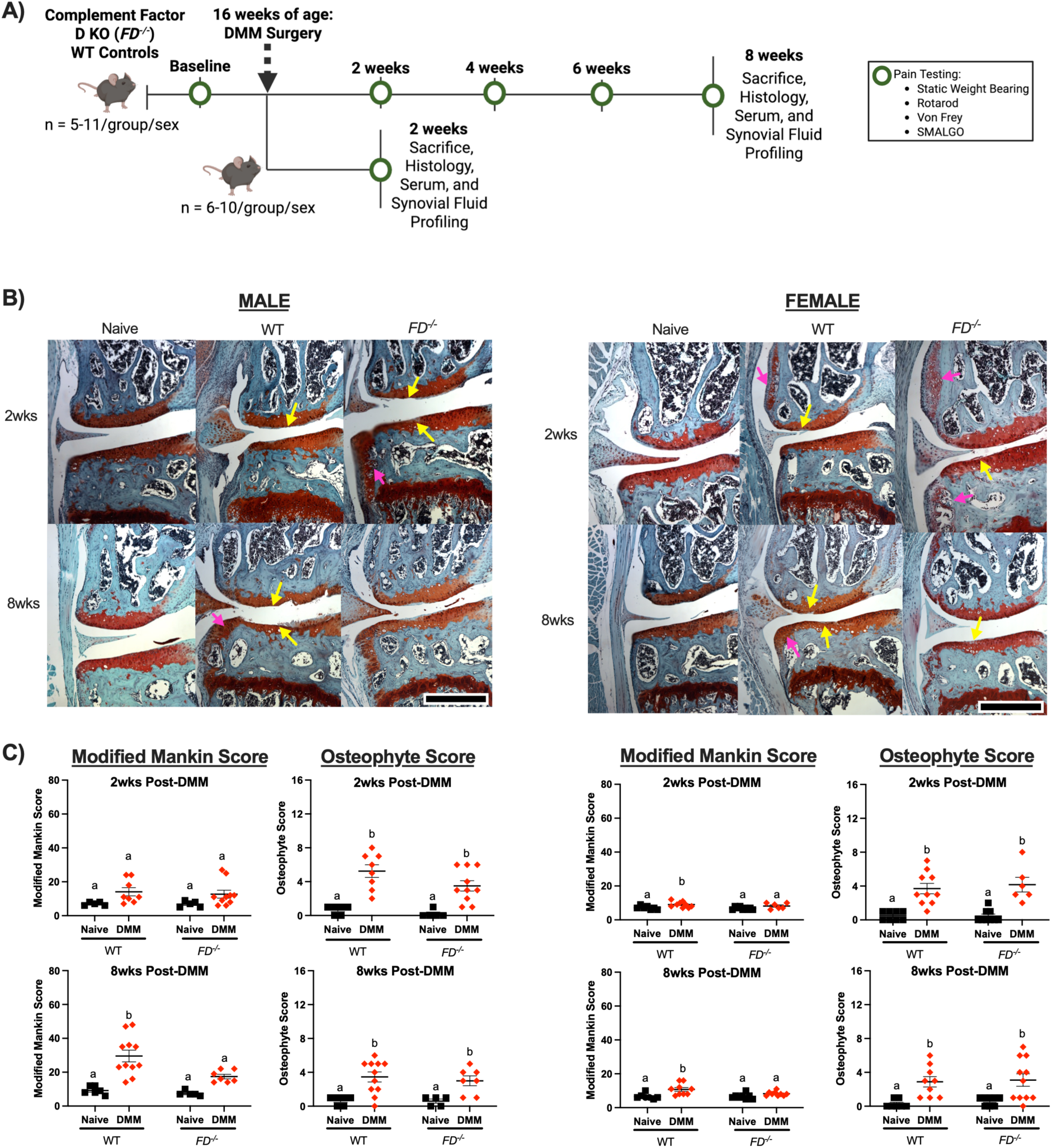
Male and female *FD^−/−^*mice displayed cartilage protection. (A) Experimental design. (B) Histological slides of the medial aspect of the left knee joint from the highest scoring joints. Sections were stained with Safranin-O/Fast Green and cartilage damage (yellow arrows) and osteophyte formation (pink arrows) are indicated. (C) Modified Mankin scores were greater in the WT groups compared to *FD^−/−^* mice 8 weeks post-DMM in male and female mice. Both sexes had greater osteophyte scores after DMM in both timepoints. Scale bar represents 500μm. Two-way ANOVA with Sidak’s post-hoc test was used to analyze between surgery. Different letters represent p < 0.0.

### Knee Joint Assessment

Knee joints were prepared according to previous methods^7,11^. In brief, joints were fixed in 4% paraformaldehyde for 24 hours and stored in 70% ethanol at 4C°. After analyzing bone microstructure, joints were decalcified in 10% formalin solution (Cal-Ex II) for 10 hours before being processed and embedded in paraffin wax. Knee joint sections were cut at 5μm thickness in the frontal plane and stained with Safranin-O/Fast Green or Hematoxylin and Eosin (H&E). Histological assessment was performed using Modified Mankin score, Osteophyte score, and Synovitis score as previously described^7,11^.

### Bone Microstructure Analysis

Whole knee joints were scanned by micro-computed tomography (SCANCO μCT50) and imaged according to the guidelines for μCT of rodent bone (energy = 55 kVP, intensity = 114 mA, 6 μm nominal voxel size, integration time = 900ms)^32^ at the Skeletal Biology and Biomechanics Core at University of California, San Francisco. Analysis of the trabecular bone in the medial and lateral tibial plateau was performed by manually contouring 2D transverse slices in the region between the growth plate and the subchondral bone. Trabecular bone volume fraction (BV/TV) and bone mineral density (BMD) were determined using the manufacturer’s analysis software. Subchondral bone was assessed similarly to exclude trabecular bone in the region between the distal surface of the femoral condyles and the growth plate. Subchondral bone thickness was determined using Fiji software as previously described^33–35^.

### Immunohistochemistry to Quantify Sensory and Sympathetic Neurites

To determine the presence of and changes in neurites in the knee joint, immunohistochemistry of calcitonin gene-related peptide (CGRP) positive and tyrosine hydroxylase (TH) positive neurons was performed as described^36^. Briefly, knee joints were embedded in paraffin wax and cut in 20μm sections at the frontal plane. Sections were blocked in 10% donkey serum and Triton-X buffer before incubation with anti-CGRP and TH primary antibodies (Bio-Rad, Millipore Sigma) overnight at 4°C. Following 3 washes of TNT buffer, sections were incubated in Alexa Fluor 647^®^ and Alexa Fluor^®^ 594 secondary antibodies (Jackson ImmunoResearch) for three hours at room temperature. The sections were then washed three times before incubation in DAPI (Sigma Aldrich) for 5 minutes. Three final washes were done before mounting with Invitrogen Fluoromount-G Mounting Medium (Fisher Scientific). Serial tiled images were taken on 10x objective images using confocal microscopy (Leica DMi8 Inverted Microscope). The region of interest was defined as the area between the synovial lining at the medial femoral quadrant, medial tibial quadrant, lateral femoral quadrant, and lateral tibial quadrant. Images were processed in Fiji to exclude bone marrow. SNT for neuroanatomy plug-in was used to quantify positive signals by manual tracing^33,37^. Total pixels of positive CGRP and TH signal were reported.

### Pain Assessments and Behavioral Testing

Pain assessments were conducted one week before DMM and 2-, 4-, 6-, and 8-weeks post-surgery in the order from least to most invasive at the same time of day. All mice were acclimatized to the behavioral suite and equipment prior to testing. Pressure-pain hyperalgesia was measured using a Small Animal Algometer^7,11^ (SMALGO, Bioseb). Three to five trials of the surgical limb and nonsurgical limb was collected by applying a steady force to the lateral aspect of each limb until the mouse showed signs of discomfort such as squeaking, paw withdrawal, or grimacing. The average of these trials was reported and a maximum value of 450g was employed to avoid tissue damage to the joint^7^.

Side-to-side limb loading was measured via static incapacitance (Bioseb). Mice were placed in the restrainer to acclimate for ∼2 minutes. Load bearing measurements for each limb were taken once the mouse was calm, had both feet on each of the force sensors, and had its paws placed on the ramp at the front. Incapacitance is measured as the difference between the right and left limbs. Three to five trials were measured, and the average was reported.

To assess mechanical allodynia, a Von Frey assay was used as described^38,39^. Mice were placed in a box with a wire mesh bottom and left to acclimate for 20 minutes. Once acclimated, force was applied to the mid-plantar of the surgical limb paw three times using one in a series of five Von Frey filaments ranging from 0.16g to 1.4g. Paw withdrawal was noted as either “positive” or “negative response”. This was then repeated with each of the five Von Frey filaments, with one repeating filament for a total of 6. The order of filaments was random between observers. Paw withdrawal patterns were assessed as previously described to determine the average 50% paw withdrawal threshold from the three repetitions^38^.

To assess motor coordination, mice were placed on a rotarod wheel with an initial speed of 4 rpm. Using the ramp function, the speed increased to a maximum of 40 rpm in 120 seconds. The time and the maximum speed at which the mouse fell were noted. The average of three to five trials was reported^40^.

### Eicosanoid Profiling by Targeted Lipidomics

Serum and synovial fluid^41^ samples were collected and extracted from male DMM mice to determine systemic and local changes in eicosanoid profiles using a novel dual extraction method that separates proteins from metabolites from a single sample^11^. In brief, all samples were extracted with methanol, vortexed, and placed at −20°C for 30 minutes to promote protein precipitation. Next, the supernatant containing metabolites was collected and dried via vacuum concentration. A 5 μL aliquot of each sample was subjected to targeted metabolomic panel of 40 known eicosanoids, analyzed at the Quantitative Metabolite Analysis Center at the University of California, San Francisco^25^ on a Shimadzu 30-AD UPLC in series with a SCIEX 7500 Triple Quadrupole Mass Spectrometer. Analytes were chromatographically separated using a Kinetex 2.6 µm Polar C18 100Å, liquid chromatography (LC) column 100 x 3.0 mm (Phenomenex, cat #00D-4759-Y0) with a mobile phase scheme of [A] water + 0.1% formic acid and [B] methanol + 0.1% formic acid. The LC method was set to a constant flow rate of 500 μL/min, and the timed linear gradient program consisted of: time=0 min, 0.10% B, time=0.1 min, 45% B, time=2 min, 45% B, time=16.5 min, 80% B, time=16.6 min, 98 % B, time=18.5 min, 98% B, time=18.6 min, 10% B, and time=20.5, 10% B. Data was collected using polarity switching with the following source parameters: CUR = 40, GS1 =60, GS2 = 70, Temp = 350 °C, ISVF = 4500V (negative mode) and 5500V (positive mode). Optimized multiple-reaction monitoring (MRM) pairs detailed in (Supplementary Tables 1 and 2) were used with a total cycle time of 0.937 seconds, dwell time of 1 ms, settling time = 15 ms, and pause time of 5.007ms. This method was based off of SCIEX’s comprehensive targeted method for lipid mediator analysis application note. Raw data was processed using a built-in SCIEX OS software package (version 2.1.6.59781) for peak picking, alignment, and quantitation.

### Statistical Analysis

All results are reported as mean ± standard error of the mean. Results from pain and behavior assays were analyzed with three-way analysis of variance (ANOVA) stratified by time point, surgery, and strain. A separate two-way ANOVA with Tukey’s post-hoc test was run by genotype and limb to determine differences within each time point. Nonparametric Spearman’s correlations were calculated to determine correlative relationships between pressure-pain threshold and osteophyte score, synovitis score, or significant eicosanoids as identified above. Lipidomic data was analyzed using an unpaired t-test on normalized abundances between WT and *FD^−/−^*mice. All other outcomes are evaluated using two-way ANOVA with Sidak’s post hoc test with strain and surgery as main effects. Heat maps and PLS-DA plots of lipidomic data were created using MetaboAnalyst. Statistical significance is defined as p < 0.05. Statistical analyses were performed using GraphPad Prism version 10.0.0 for Mac (GraphPad Software, Boston, Massachusetts USA, www.graphpad.com).

## RESULTS

### Male and female *FD^−/−^* mice displayed cartilage protection after DMM surgery

As expected, *FD^−/−^* male and female mice were protected from DMM-induced structural damage as shown in images of the highest Mankin Scores in each group (Figure 1b). Concordant with previous analysis at 12-weeks post-DMM^11^, male WT mice demonstrated significantly greater Modified Mankin scores between DMM and naïve groups at 8 weeks post-DMM (p<0.001), *FD^−/−^* mice displayed no significant differences between DMM and naïve groups (Figure 1c). Similarly, female WT mice exhibited greater Modified Mankin scores at both 2 and 8 weeks (p=0.01, p=0.001), although there were no cartilage changes in *FD^−/−^* mice. Despite cartilage protection in *FD^−/−^* mice, significant osteophyte formation was present in both strains. WT and *FD^−/−^*DMM groups of male and female mice had greater osteophyte scores compared to the naïve controls at 2 weeks (male: p<0.001, p<0.01; female: p<0.001, p<0.0001) and 8 weeks (male: p<0.01, p=0.02; female: p=0.01, p<0.01) post-DMM (Figure 1c).

Similarly, synovitis scores were greater in all DMM groups compared to the naïve control at both 2 weeks (male: WT p<0.01, *FD^−/−^*p<0.001; female: WT p<0.001, *FD^−/−^* p<0.001) and 8 weeks (male: WT p<0.01, *FD^−/−^*<0.001; female: WT p<0.001, *FD^−/−^*p<0.001) post-DMM (Figure 2). Interestingly, *FD^−/−^* presented worsening synovitis scores compared to WT at 2 weeks post-DMM (p=0.02) (Figure 2).

**Figure 2.**
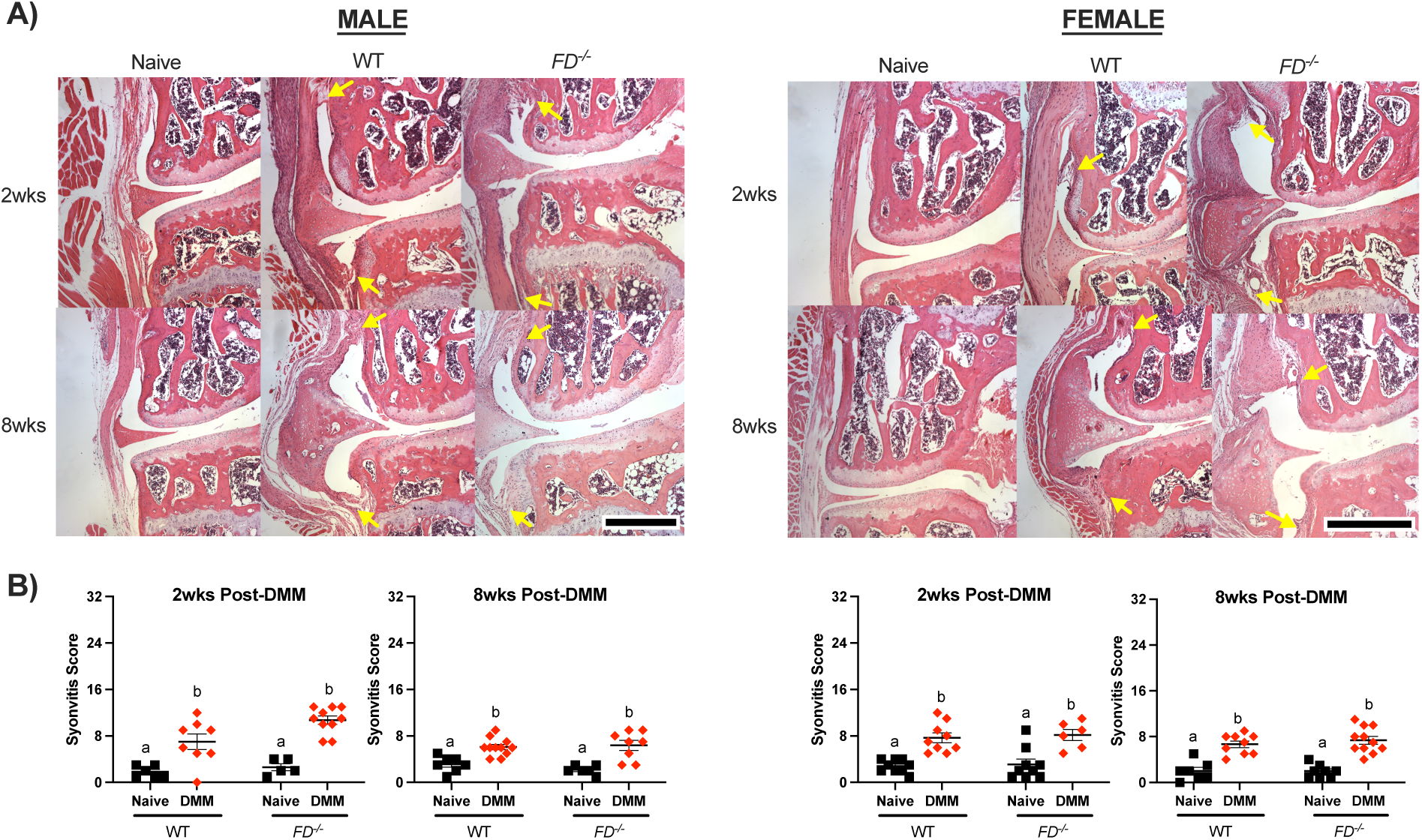
Synovitis scores increased after DMM. (A) Representative images of histological slides stained with Hematoxylin and Eosin (H&E). Yellow arrows indicate increased thickening, cellularity, and fibrosis of the synovial lining. (B) Although all DMM groups scored higher for synovitis compared to naïve, male *FD^−/−^* mice displayed greater scores than the WT DMM group at 2 weeks post-DMM. Scale bar represents 500μm. Two-way ANOVA with Sidak’s post-hoc test was used to analyze between surgery. Different letters represent p < 0.05.

Analysis of bone changes revealed subchondral bone thickening in the lateral posterior compartment in WT (p=0.01), but not *FD^−/−^* male at 8 weeks post-DMM (Figure 3). There were no significant differences in subchondral thickness at 2 weeks post-DMM, or in female mice. There were no differences in BV/TV of the proximal tibial epiphysis due to surgery, strain, or sex (Supplementary Figure 1).

**Figure 3.**
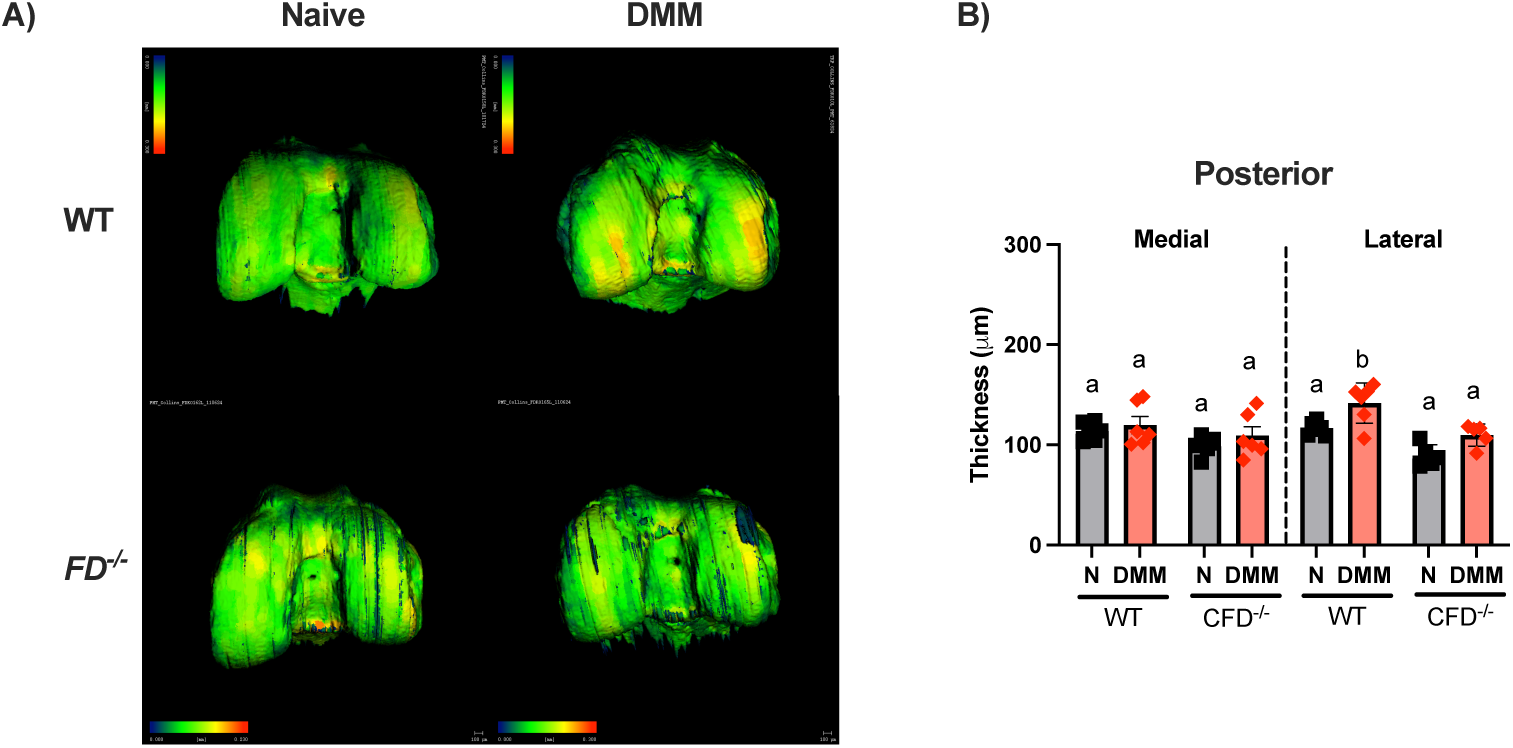
MicroCT analysis revealed subchondral bone thickening 8 weeks post-DMM in male mice. (A) Representative images of the posterior distal femur. (B) Subchondral bone thickness of the distal femur was greater in the posterior lateral compartment in WT mice. Scale bar represents 100μm. Two-way ANOVA with Sidak’s post-hoc test was used to analyze between surgery. Different letters represent p < 0.05.

Preliminary analysis of a subset of joints to quantify CGRP+ and TH+ neurites in the joint did not demonstrate any clear changes between surgical groups, strain, or sex in a subset of animals (Supplementary Figure 2).

### Pressure-pain Hyperalgesia was increased in *FD^−/−^* mice at 2-weeks, and persisted to 8 weeks

Pain and behavior assays were conducted one week before DMM at baseline, and 2, 4, 6, and 8 weeks after surgery. The surgical limb of WT and *FD^−/−^*DMM male mice displayed decreases in pressure-pain threshold as early as 2 weeks and through 8 weeks post-DMM when compared to their contralateral limb (WT: p= <0.001, p= 0.03, p= 0.02, p <0.001; *FD^−/−^*: p<0.002, p= 0.01, p <0.01). Surprisingly, female *FD^−/−^* mice showed similar pressure-pain hyperalgesia to male *FD^−/−^* and WT mice starting from 2 weeks throughout 8 weeks post-DMM (p<0.001 p<0.01, p<0.01, p<0.01) but WT female mice did not display significantly decreased thresholds compared to the contralateral limb until 6 weeks post-DMM, which persisted to 8-weeks (p= 0.03, p<0.01). There were no differences in pressure-pain thresholds between *FD^−/−^* and WT DMM groups when evaluated within time points (Figure 4a) in male mice. Male mice exhibited decreased loading of the DMM limb following injury in both *FD^−/−^* and WT mice. However, while WT maintained offloading consistently through 6 weeks (WT: p<0.001, p <0.05, p<0.01), *FD*^−/−^ only demonstrated significantly increased DMM limb offloading at 2 weeks post-DMM compared to naïve mice (p<0.001). In contrast, female mice displayed more modest DMM limb offloading behavior. Only female *FD^−/−^* mice exhibited significantly different side-to-side limb loading acutely 2 weeks after DMM, and again chronically at 6 and 8 weeks post-DMM compared to naïve (p=0.03, p< 0.01) (Figure 4b). There were no differences in mechanical allodynia (Figure 4c) or motor coordination (Figure 4d) between DMM and naïve groups, although male *FD^−/−^* DMM mice demonstrated significantly longer time on the rotarod than the WT group at later timepoints (p=0.02; p=0.02).

**Figure 4.**
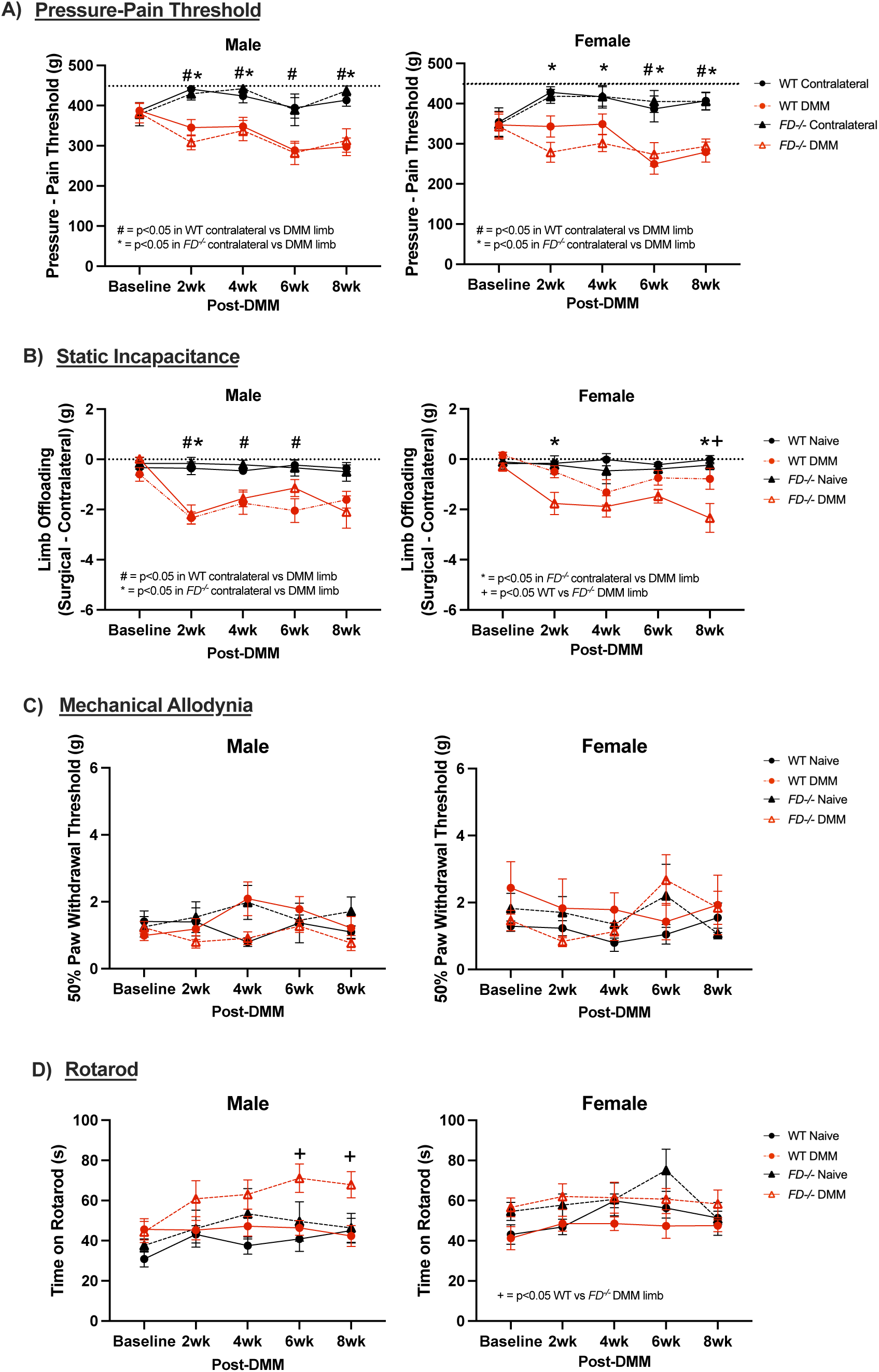
Pain phenotypes appeared at 2 weeks post-DMM and persisted through 8 weeks in both strains. (A) Male mice displayed clear pressure-pain hyperalgesia at 2 weeks and maintained for an 8-week observation period. Female *FD^−/−^* mice exhibited hyperalgesia at each timepoint. There were no differences between *FD^−/−^*and WT in either sexes. (B) Offloading of the surgical limb after DMM was present in both male and female mice. (C) Mechanical allodynia and (D) motor coordination was no different between groups. Two-way ANOVA analysis with Tukey’s post-hoc was done at each timepoint with surgery and strain as main effects. # denotes p<0.05 when comparing contralateral and DMM limb in WT mice, * denotes p<0.05 when comparing contralateral and DMM limb in *FD^−/−^* mice, + denotes p<0.05 when comparing WT and *FD^−/−^* DMM mice.

To understand if the increased pain phenotypes in *FD^−/−^*mice were explained by changes in histology, correlation analysis was performed with pressure-pain hyperalgesia and static incapacitance with osteophyte and synovitis scores. Pressure-pain threshold was significantly correlated with osteophyte scores in WT male mice at 8 weeks post-DMM (ρ = −0.72; p= 0.03) (Supplementary Figure 3a), although there were no differences detected in sensory or sympathetic neurites in the osteophyte region in the preliminary set of images evaluated (Supplementary Figure 2). Additionally, offloading of the surgical limb was significantly correlated with osteophyte scores in WT female mice at 2 weeks post-DMM (ρ = −0.66; p = 0.04). However, no other significant correlations between pressure-pain threshold or static incapacitance and osteophytes were observed in male or female *FD^−/−^* mice (Supplementary Figure 3b). In comparison, synovitis scores were significantly correlated with pressure-pain threshold in WT females at 8 weeks post-DMM (ρ = 0.775; p = 0.05) (Supplementary Figure 4a). However, there were no significant correlations in any other WT or *FD^−/−^* mice between pressure-pain threshold or static incapacitance with synovitis of both sexes. (Supplementary Figure 4).

### Early eicosanoid changes in *FD^−/−^* mice were detected in synovial fluid and serum

To determine the role of known eicosanoids – fat-secreted bioactive lipid mediators – in the *FD^−/−^* pain phenotype, lipidomic profiles were assessed from serum and synovial fluid of the surgical limb in DMM male mice at 2 and 8 weeks post-DMM using LC-MS. PLS-DA plots demonstrate a clear separation of lipidomic profiles at 2 weeks post-DMM in the serum and synovial fluid. *FD^−/−^*mice 2 weeks post-DMM displayed significant decreases of wound healing factor 12-HHT(12-hydroxyheptadecatrienoic acid)^42^ (p<0.01) in the serum, and pain suppression factors 14,15-EET (14,15-epoxyeicosatrienoic acid) and 10,11-EpDPA (10, 11-docosahexaenoic acid)^43–46^ (p=0.02, p=0.03) in the synovial fluid when compared to WT (Figure 5a).

**Figure 5.**
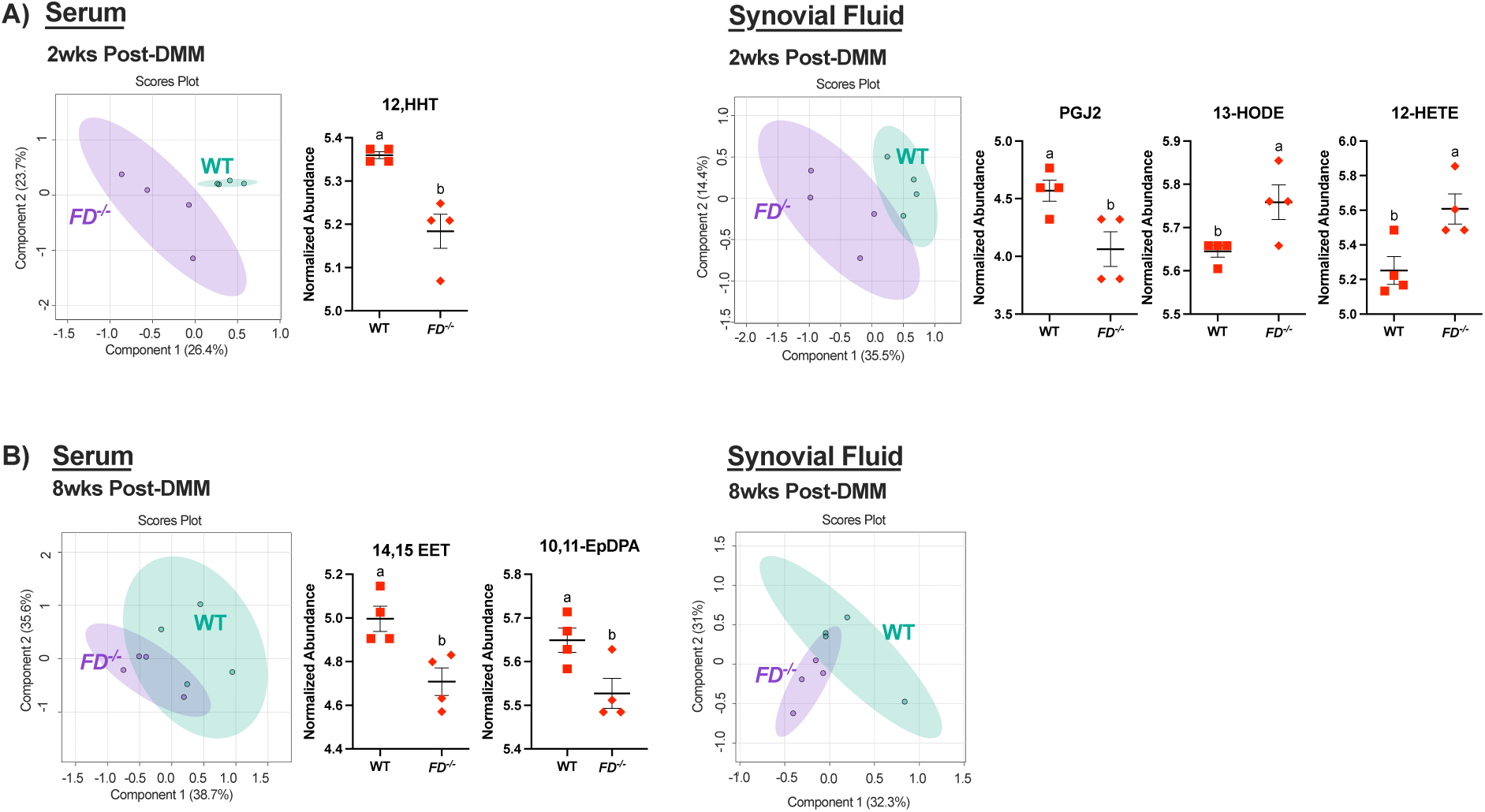
*FD^−/−^* mice exhibited dysregulation of lipid profiles in the serum and synovial fluid. (A) PLS-DA plots show separation in systemic and synovial fluid lipidomic profiles of male *FD^−/−^* compared to WT at 2 weeks post-DMM. Wound-healing factor (12-HHT) is lower in the serum of FD^−/−^ in comparison to WT. Pain suppression factors (PGJ2, 13-HODE, 12-HETE) are lower in the synovial fluid of the surgical limb in *FD^−/−^* mice compared to WT. (B) At 8 weeks post-DMM, more overlap is observed in serum and SF. Differences in factors related to pain (14,15-EET, 10,11-EpDPA) are found in the serum, not the synovial fluid, when comparing *FD^−/−^* to WT mice. Unpaired t-test was used to analyze between strains. Different letters represent p < 0.05.

### Systemic eicosanoid changes in *FD^−/−^* mice were distinct at 8-weeks post DMM

At 8 weeks, serum and synovial fluid eicosanoid profiles showed moderate overlap as shown in the PLS-DA plots (Figure 5b). *FD^−/−^* mice displayed decreases in pain suppression factor (prostaglandin J2, or PGJ2)^47^ (p=0.03) and increases in pain driving factors 12-HETE (12-hydroxyeicosatetraenoic acid) and 13-HODE (13-hydroxyoctadecadienoic acid)^48,49^ (p=0.02, p=0.04) in the serum compared to WT. However, there were no differences in the eicosanoid profiles in the synovial fluid at 8 weeks post-DMM (Figure 5b).

To demonstrate the differential eicosanoid profiles in each group and over time, heat maps and PLS-DA plots of four-group comparisons between strain and time point are presented in Supplementary Figures 5 and 6. Of note, there were no significant correlations between normalized eicosanoid abundance and pressure-pain threshold or static incapacitance in combined populations of *FD^−/−^*and WT in the serum or synovial fluid at 2 or 8 weeks post-DMM (Supplementary Figure 7).

## DISCUSSION

The assumption that all pain in OA is derived from peripheral structural insults or changes is a major barrier to understanding the mechanisms of OA pain. This study leverages a novel preclinical mouse model that we have recently demonstrated recapitulates the clinical reports of discordance between pain and structural damage in knee OA^11^. Through constitutive knockout of *Cfd*, we establish that FD is a key mediator in driving cartilage damage in the onset of knee OA as early as 2 weeks post-DMM in male and female mice. Despite cartilage protection, *FD^−/−^* animals challenged with DMM exhibited pain phenotypes that were not simply explained by histological assessments of osteophytes or synovitis, or neurite growth in the joint. These data indicate that structural damage in the joint is not the primary driver for pain in acute and early OA in the *FD^−/−^* model. Analysis of pain outcomes 2 weeks post-DMM demonstrated heightened pain sensitivity due to DMM that remained consistent throughout early onset of OA in both groups of male mice. Surprisingly, we observed that with loss of FD, male and female mice demonstrated similar pressure-pain hyperalgesia and static incapacitance, which differed dimorphically in male and female WT mice. This suggests that FD may contribute to sexual dimorphisms in the development of pain with DMM and as OA becomes entrenched. Lastly, we used an innovative targeted approach to map the role of eicosanoids in synovial fluid and serum over time, which corroborates the notion that eicosanoids may be downstream mediators of FD-driven pain in OA in mice. We discovered that loss of FD in male mice resulted in increases in pain-driving (12-HETE, 13-HODE)^48,49^ and decreases in pain-suppressive associated (PGJ2, 14,15-EET, and 10,11-EpDPA) ^43–47^ eicosanoids after DMM when compared to WT, thus these eicosanoids may be novel targets for the development of drugs to target OA pain.

Here, we demonstrate that FD drives cartilage changes in OA in male and female mice. Congruent with existing literature, *FD^−/−^* mice were protected from cartilage damage compared to WT after DMM. Recent studies, including our own, indicate that male *FD^−/−^* mice showed significantly less cartilage damage in spontaneous and DMM models of established OA with similar levels of pressure-pain hyperalgesia 12 weeks post-DMM^9–11^. However, previous studies focused on male mice, so little is known about the FD and alternative complement signaling in female mice. Because female mice have been reported to display more modest cartilage damage after DMM^50^, they are often excluded from OA preclinical studies. Interestingly, the present data provide evidence to the contrary. With sufficient sample sizes, WT female mice displayed statistically significant cartilage damage that was not present in the *FD^−/−^* group, indicating that the chondroprotective effect of FD is not sex-dependent. These data are important for two key reasons. First, this supports the notion that female mice challenged with DMM is an intriguing model of OA pain. Second, our findings indicate that there may be sexual dimorphisms in the mechanism of FD-mediated pain-structure discordance that should be considered. WT female mice demonstrated more pain for a given amount of cartilage damage, which is concordant with reports of increased pain severity in female patients^12^, and supports the notion that holistically phenotyping preclinical mouse studies with pain and behavior may provide more translationally relevant targets and mechanistic understanding.

Our findings illustrate that early pressure-pain hyperalgesia and static incapacitance are the result of DMM injury. Consistent with previous studies^51^, pressure-pain hyperalgesia thresholds for both WT and *FD^−/−^* mice significantly decrease at two weeks in male mice and remained consistent through the 8-week observation period, congruent with our previous work^11^. In females, to our surprise, *FD^−/−^* mice demonstrated a similar decrease as males in pressure-pain threshold at 2 weeks that was maintained through 8 weeks post-DMM when compared to the contralateral limb. While we did not observe changes in mechanical allodynia in this study due to DMM or loss of FD, other studies have reported that allodynia thresholds were lowest at 2-8 weeks post-DMM before a reversal of sensitivity after 8 weeks^14,52^. Taken together, these findings indicate comparable pain response between *FD^−/−^* male and female mice despite female mice exhibiting less severe cartilage damage, and as such, we reject the initial hypothesis that *FD^−/−^* male mice would have a more profound pain phenotype compared to female mice.

While several studies demonstrate a role for FD and alternative complement signaling in driving cartilage damage, FD’s influence on pathological changes in other joint changes after DMM is unclear. This is particularly important because it is known that complement factors are produced and activated in almost all joint tissues^53^, including the infrapatellar fat pad, which we recently showed strong FD expression in adiponectin-positive barcodes using spatial transcriptomics^11^. Furthermore, while there is a lack of studies that directly investigate the role of FD on osteophyte formation in OA, we have previously reported a robust osteophyte phenotype in *FD^−/−^* mice 12 weeks post DMM^11^. Additionally, several studies have reported the absence of FD inhibits synovitis^9,10^. In this present study, we observe synovitis and osteophytes coincident with cartilage protection, similar to previous work in fat-free lipodystrophic mice^7^ and 12-week evaluation of *FD^−/−^* mice^11^. These results highlight the potential separable nature of joint tissues in OA. Multiple, perhaps contradictory, roles for FD in response to DMM injury and the manifestation of pain require further investigation to understand its multiple key roles in OA progression and pathological joint tissue changes.

Although osteophytes and synovitis have been implicated in pain in clinical and preclinical models^54–57^, our findings indicate that the role of FD on pain is not simply explained by semiquantitative measured changes in these outcomes by histology. Pressure-pain threshold was not significantly correlated to osteophyte or synovitis scores, suggesting that structural changes of the knee cannot fully explain increased pain sensitivity in either sex. However, these measures are crude, and many more advanced techniques and immune profiling are available. It is important to note that, by design, our work focuses on early onset OA to capture responses to injury (acute) and how FD affects the joint and pain over time (chronic). To date, most preclinical and clinical studies investigate the relationship between pain and structure during established and severe OA progression^11,55^. Precise and focused work is needed to confirm or reject the direct role of osteophytes and synovitis on *FD^−/−^* driven pain, which remains a very promising avenue for future work. Preliminary analysis of CGRP+ and TH+ nerve endings did not reveal any differences between strain, surgery, or sex, although greater sample sizes are needed to determine whether FD plays a role in differential sensory or sympathetic neurite sprouting after DMM. Altogether, these findings demonstrate that pain sensitivity is not simply explained by these structural changes in early-onset OA. Rather, this suggests that pain may be regulated elsewhere in, or potentially outside, of the joint. Twelve weeks after DMM, we observed alterations in pathways associated with neutrophil trap formation, immunodeficiency, insulin signaling, and calcium signaling in bulk RNA sequencing of the dorsal root ganglia neurons that innervate the knee of *FD^−/−^* vs WT mice^11^. Our ongoing efforts aim to profile the DRGs from 2 weeks and 8 weeks post-DMM to uncover acute and chronic changes to these nerves and how they may help explain the early and lasting pain phenotype in the present *FD^−/−^* male and female mice.

Eicosanoids can be triggered by leptin in the synovium^58^ and contribute to the perception of pain through cytokine and COX2 activity^28,29,31,58^. As such, eicosanoids are the targets in NSAID treatment, such as celecoxib, for OA pain. Interestingly, celecoxib has known sexually dimorphic responses in treatment efficacy. Male Sprague-Dawley rats demonstrate better pain relief compared to females^59^. Because NSAIDS can modulate the complement system^24^, we utilized a targeted lipidomics panel to understand how eicosanoids are mediated by FD in response to injury and OA disease entrenchment. Moderate overlap of eicosanoid profiles in serum and synovial fluid for both time points suggests that eicosanoid changes occur in both serum and synovial fluid but in a time-dependent manner. As expected, PLS-DA plots show distinct clusters at 2 weeks but more overlap at 8 weeks due to FD’s role in activating the alternative pathway in response to tissue injury^60^. Furthermore, lower levels of wound-healing factor 12-HHT were measured in *FD^−/−^* compared to WT after DMM. These findings suggest that FD likely plays a bigger role in the acute inflammatory phase following DMM. Differences were detected in the serum and synovial fluid at this early time point, highlighting both the local and systemic effects of FD after DMM and further supporting the paradigm that OA is a systemic disease that affects the rest of the body, and can be affected by tissues outside the joint^7,11^. Strikingly, eicosanoid profiling demonstrated that loss of FD results in the downregulation of pain suppression and upregulation of pain-driving factors that may be acting synergistically and explain the strong pain phenotype measured in these studies. The combined effects of reducing inhibitory factors and additive effects of pain driving factors are concordant with the pronounced pain phenotype in *FD^−/−^*. However, because pain levels were similar between *FD^−/−^* male and female mice, we focused on analyzing eicosanoid profiles in male mice. Our future work will investigate differences in the targeted eicosanoid profile and untargeted metabolome of female *FD^−/−^*and WT mice to better gain insight into how FD may be influencing pain in OA.

2 weeks after DMM surgery, we observed lower levels of PGJ2, an antagonist of the inflammatory mediator prostaglandin E2 (PGE2)^61^, in the synovial fluid of the surgical limb in *FD^−/−^* mice compared to WT. Intra-muscular injection of PGJ2 has been shown to target both PPARγ and opioid receptors to prevent muscle hyperalgesia in rats^47^. In contrast, levels of 12-HETE, a molecule shown to amplify PGE2 signaling^49^, and 13-HODE, an endogenous TRPV1 ligand implicated in inflammatory pain^48,62^, were higher in the synovial fluid of *FD^−/−^*mice at 2 weeks post-DMM when compared to WT. 13-HODE is implicated in inflammatory pain, and administration of anti-13-HODE antibodies has been shown to significantly reduce inflammatory hyperalgesia^62^. At 8 weeks post-DMM, lower levels of 14,15-EET and 10,11-EpDPA were detected in *FD^−/^*^−^ mice compared to WT. Both eicosanoids are associated with anti-nociceptive effects in murine models and have been shown to reduce to neuropathic and inflammatory pain when injected^43–46^. Furthermore, lower levels of EETs have been associated with chronic joint pain in OA in both human and mouse models^45^. In comparison, while there were no differences in synovial fluid eicosanoid profiles, clear changes in pain related factors in the serum were detected at 8 weeks.

Although we did not find any significant correlations between pressure-pain threshold and the eicosanoid factors identified above, this may be due to: (1) the multiple functions FD may serve in OA pathogenesis, (2) insufficient sample size to identify if a relationship exists with these eicosanoid factors and OA outcomes, and (3) that the 40 eicosanoids measured do not completely capture the eicosanoid profiles of FD and WT mice after DMM. Thus, our future work will leverage untargeted metabolomics approaches to provide insight into other mediators that play a significant role in FD-associated pain and uncover novel molecular signatures to identify novel pain biomarkers and therapeutic targets for OA.

While this study provides new knowledge on the mechanisms of pain and structure mediated by FD post injury and over time in both male and female mice, it is not without limitations. First, the analysis of changes in nerve endings within the joint was preliminary. Additional samples are needed to thoroughly characterize whether FD plays a role in neurite sprouting at the joint. Second, we focused on male DMM samples for eicosanoid phenotyping. Therefore, future work will consider both sexes. Third, while mouse models are valuable in disentangling mechanisms of OA pain and structure, there are immunological differences between mice and humans, specifically in alternative complement vs classical complement signaling^63–66^. Future work will be needed in humanized mice to validate the role of FD driving pain as a key next step in translating these findings to the clinic and developing the eicosanoid targets.

In conclusion, this study contributes to the growing body of evidence that OA is a systemic disease that can be driven by factors outside the joint and can reciprocally affect the whole body. We demonstrate that valuable mechanistic information on OA pain can be gleaned by leveraging models that exhibit structure-pain discordance in response to DMM and by profiling this discordance over time. By global deletion of FD, we are able to recapitulate clinical pain-structure discordance, which allowed us to characterize the time course of pain and cartilage damage separately. We determined that *FD^−/−^* has a chondroprotective effect and influences pressure-pain sensitivity in early onset OA in both male and female mice. Furthermore, our findings suggest that *FD^−/−^* may contribute to sexual dimorphisms in pressure-pain hyperalgesia and side-to-side limb loading in WT animals. Targeted eicosanoid profiling of 40 known metabolites^25^ revealed changes in pain regulation both locally and systemically and demonstrates its potential in identifying key mechanistic insights into the relationship between fat, OA, and pain. These findings will lay the groundwork for identifying new fat-derived therapeutic targets downstream of FD that will be able to address pain and structure in OA pathogenesis.

## DECLARATIONS

## Ethics approval and consent to participate

All experimental procedures were approved by the University of California, San Francisco Institutional Animal Care and Use Committee.

## Consent for publication

Not applicable.

## Acknowledgements

We would like to thank the Skeletal Biology and Biomechanics Core at UCSF Core Center for Musculoskeletal Biology and Medicine for their technical support with microCT scanning and to the UCSF Quantitative Metabolite Analysis Center (QMAC) for their support in conducting lipidomic analysis. QMAC is made possible by support from the Benioff Center for Microbiome Medicine, ImmunoX, and the Program for Breakthrough Biomedical Research (PBBR). We would like to extend additional gratitude to John Atkinson and Xiaobo Wu for the *FD^−/−^*mice used in this study and for scientific discussions. We would also like to thank Christine Pham for scientific discussions. *FD^−/−^* mice are available from J Atkinson and X. Wu from Washington University in St. Louis by MTA or reasonable request. Lastly, the authors would also like to thank Kristen W.Y. Chan and Reyna E. Villa for their histological analysis support for this project.

## AUTHOR CONTRIBUTIONS

P.T. and K.C. conceived and executed the studies, analyzed data, and wrote the main manuscript. B.A. managed the animal colony and conducted pain testing in Figure 4. J.K. stained and imaged histology data in figures 1 and 2. M.R. imaged and analyzed nerve endings. S.S., T.K., and A.G. analyzed microCT data in figure 3. D.D. and H.W. conducted the lipidomic experiment in figure 5. H.W. additionally analyzed and interpreted lipidomic data. All authors reviewed the manuscript.

