## Supplemental Information for "Complement factor D (adipsin) mediates pressure-pain hypersensitivity post destabilization of medial meniscus injury"

**This PDF file includes:**

Supplementary Tables S1-S2
Supplementary Figures S1-S7

**Supplemental Information**

**Supplementary Table 1. Negative Mode**

| **Compound ID** | **Q1** | **Q3** | **Edit Dwell** | **Dwell Time** | **EP** | **CE** | **CXP** | **Expected RT (min)** | **RT Tolerance (s)** |
| --- | --- | --- | --- | --- | --- | --- | --- | --- | --- |
| PGE2-d4 (355/193) | 355.2 | 193.1 | 1 | 1 | 10 | 25 | 12 | 8.1 | 30 |
| 15-deoxy-PGJ2-d4 (319/275) | 319.2 | 275.1 | 1 | 1 | 10 | 20 | 12 | 13.6 | 30 |
| 8-iso-PGF2a-d4 (357/197) | 357.2 | 197.1 | 1 | 1 | 10 | 30 | 12 | 7.2 | 30 |
| LTB4-d4 (339/197) | 339.2 | 197.1 | 1 | 1 | 10 | 22 | 12 | 12.1 | 30 |
| LXA4-d5 (356/115) | 356.2 | 115.1 | 1 | 1 | 10 | 19 | 12 | 8.9 | 30 |
| 11,12-EET-d11 (327/171) | 327.2 | 171.1 | 1 | 1 | 10 | 20 | 12 | 16.5 | 30 |
| 5-HETE-d8 (327/116) | 327.2 | 116.1 | 1 | 1 | 10 | 17 | 12 | 15.9 | 30 |
| 12-HETE-d8 (327/184) | 327.2 | 184.1 | 1 | 1 | 10 | 20 | 12 | 15.6 | 30 |
| 15-HETE-d8 (327/226) | 327.2 | 226.1 | 1 | 1 | 10 | 20 | 12 | 15.3 | 30 |
| RvE1-d4 (353/197) | 353.2 | 197.1 | 1 | 1 | 10 | 22 | 12 | 5.9 | 30 |
| RvD2-d5 (380/141) | 380.2 | 141.1 | 1 | 1 | 10 | 23 | 12 | 8.7 | 30 |
| RvD3-d5 (380/152) | 380.2 | 152.1 | 1 | 1 | 10 | 25 | 12 | 8.6 | 30 |
| Maresin 1-d5 (364/221) | 364.2 | 221.1 | 1 | 1 | 10 | 20 | 12 | 12.1 | 30 |
| 6k-PGF1a (369/163) | 369.2 | 163.1 | 1 | 1 | 10 | 35 | 12 | 6.8 | 30 |
| PGE2 and PGD2 (351/189) | 351.2 | 189.1 | 1 | 1 | 10 | 25 | 12 | 8.15 | 30 |
| PGE2 (351/175) | 351.2 | 175.1 | 1 | 1 | 10 | 25 | 12 | 8.06 | 30 |
| 15-keto-PGE2 (349/113) | 349.2 | 113.1 | 1 | 1 | 10 | 28 | 12 | 7.4 | 30 |
| 13,14-dihydro-15-keto PGE2 (351/175) | 351.2 | 175.1 | 1 | 1 | 10 | 26 | 12 | 8.1 | 30 |
| PGD2 (351/233) | 351.2 | 233.1 | 1 | 1 | 10 | 16 | 12 | 8.22 | 30 |
| 11-beta-PGF2a (353/193) | 353.2 | 193.1 | 1 | 1 | 10 | 30 | 12 | 7.5 | 30 |
| 13,14-dihydro-15-keto PGD2 (351/207) | 351.2 | 207.1 | 1 | 1 | 10 | 26 | 12 | 8.8 | 30 |
| PGJ2 (333/271) | 333.2 | 271.1 | 1 | 1 | 10 | 22 | 12 | 10.1 | 30 |
| 15-deoxy-PGJ2 (315/271) | 315.2 | 271.1 | 1 | 1 | 10 | 20 | 12 | 13.6 | 30 |
| PGF2a (353/193) | 353.2 | 193.1 | 1 | 1 | 10 | 34 | 12 | 8.55 | 30 |
| 15-keto PGF2a (351/113) | 351.2 | 113.1 | 1 | 1 | 10 | 35 | 12 | 8 | 30 |
| 13,14-dihydro-15-keto PGF2a (353/291) | 353.2 | 291.1 | 1 | 1 | 10 | 25 | 12 | 9.1 | 30 |
| 20-OH PGF2a (369/193) | 369.2 | 193 | 1 | 1 | 10 | 35 | 12 | 3.22 | 30 |
| TXB2 (369/169) | 369.2 | 169.1 | 1 | 1 | 10 | 22 | 12 | 7.7 | 30 |
| 12-HHT (279/179) | 279.2 | 179.1 | 1 | 1 | 10 | 22 | 12 | 13 | 30 |
| LTB4 (335/195) | 335.2 | 195.1 | 1 | 1 | 10 | 22 | 12 | 12.1 | 30 |
| 20-OH-LTB4 (351/195) | 351.2 | 195.1 | 1 | 1 | 10 | 24 | 12 | 6.6 | 30 |
| 20-COOH-LTB4 (365/195) | 365.1 | 195.1 | 1 | 1 | 10 | 25 | 12 | 6.16 | 30 |
| LXA4 (351/115) | 351.2 | 115.1 | 1 | 1 | 10 | 20 | 12 | 8.9 | 30 |
| 15(R)-LXA4 (351/115) | 351.2 | 115.1 | 1 | 1 | 10 | 20 | 12 | 9.1 | 30 |
| LXB4 (351/221) | 351.2 | 221.1 | 1 | 1 | 10 | 22 | 12 | 8.4 | 30 |
| 5,15-diHETE (335/115) | 335.2 | 115.1 | 1 | 1 | 10 | 22 | 12 | 11.7 | 30 |
| 5S,6R-diHETE (335/115) | 335.2 | 115.1 | 1 | 1 | 10 | 20 | 12 | 13.6 | 30 |
| 5,6 EET (319/191) | 319.2 | 191.1 | 1 | 1 | 10 | 21 | 12 | 16.7 | 30 |
| 8,9 EET (319/155) | 319.2 | 155.1 | 1 | 1 | 10 | 21 | 12 | 16.6 | 30 |
| 11,12 EET (319/167) | 319.2 | 167.1 | 1 | 1 | 10 | 21 | 12 | 16.5 | 30 |
| 14,15 EET (319/219) | 319.2 | 219.1 | 1 | 1 | 10 | 15 | 12 | 16.2 | 30 |
| 5,6 DiHETrE (337/145) | 337.2 | 145.1 | 1 | 1 | 10 | 25 | 12 | 14.4 | 30 |
| 8,9 DiHETrE (337/127) | 337.2 | 127.1 | 1 | 1 | 10 | 30 | 12 | 13.9 | 30 |
| 11,12 DiHETrE (337/167) | 337.2 | 167.1 | 1 | 1 | 10 | 25 | 12 | 13.6 | 30 |
| 14,15 DiHETrE (337/207) | 337.2 | 207.1 | 1 | 1 | 10 | 25 | 12 | 13.4 | 30 |
| 5-HETE (319/115) | 319.2 | 115.1 | 1 | 1 | 10 | 21 | 12 | 15.9 | 30 |
| 9-HETE (319/167) | 319.3 | 167.2 | 1 | 1 | 10 | 21 | 12 | 15.8 | 30 |
| 11-HETE (319/167) | 319.2 | 167.1 | 1 | 1 | 10 | 21 | 12 | 15.4 | 30 |
| 12-HETE (319/179) | 319.2 | 179.1 | 1 | 1 | 10 | 21 | 12 | 15.6 | 30 |
| 15-HETE (319/219) | 319.2 | 219.1 | 1 | 1 | 10 | 19 | 12 | 15.3 | 30 |
| 18-HETE (319/261) | 319.2 | 261.2 | 1 | 1 | 10 | 25 | 12 | 14.74 | 30 |
| 20-HETE (319/289) | 319.2 | 289.1 | 1 | 1 | 10 | 25 | 12 | 14.7 | 30 |
| 8-iso-PGF2a (353/193) | 353.2 | 193.1 | 1 | 1 | 10 | 30 | 12 | 7.2 | 30 |
| AA (303/259) | 303.2 | 259.1 | 1 | 1 | 10 | 16 | 12 | 17.8 | 30 |
| RvE1 (349/195) | 349.2 | 195.1 | 1 | 1 | 10 | 22 | 12 | 5.9 | 30 |
| RvE2 (333/199) | 333.2 | 199.1 | 1 | 1 | 10 | 22 | 12 | 10.1 | 30 |
| RvE2(333/115) | 333.2 | 115.1 | 1 | 1 | 10 | 22 | 12 | 10.1 | 30 |
| 5,15-DiHEPE (333/115) | 333.2 | 115.1 | 1 | 1 | 10 | 22 | 12 | 10.5 | 30 |
| RvE3 (333/201) | 333.2 | 201.1 | 1 | 1 | 10 | 20 | 12 | 12 | 30 |
| LXA5 (349/115) | 349.2 | 115.1 | 1 | 1 | 10 | 20 | 12 | 7.5 | 30 |
| 5-HEPE (317/115) | 317.2 | 115.1 | 1 | 1 | 10 | 19 | 12 | 14.6 | 30 |
| 11-HEPE (317/167) | 317.2 | 167.1 | 1 | 1 | 10 | 19 | 12 | 14.3 | 30 |
| 12-HEPE (317/179) | 317.2 | 179.1 | 1 | 1 | 10 | 19 | 12 | 14.4 | 30 |
| 15-HEPE (317/219) | 317.2 | 219.1 | 1 | 1 | 10 | 18 | 12 | 14.3 | 30 |
| 18-HEPE (317/259) | 317.2 | 259.1 | 1 | 1 | 10 | 16 | 12 | 14.1 | 30 |
| 8,9-EpETE (317/127) | 317.2 | 127.1 | 1 | 1 | 10 | 20 | 12 | 15.4 | 30 |
| 11,12-EpETE (317/167) | 317.2 | 167.1 | 1 | 1 | 10 | 20 | 12 | 15.4 | 30 |
| 14,15-EpETE (317/207) | 317.2 | 207.1 | 1 | 1 | 10 | 20 | 12 | 15.4 | 30 |
| 17,18-EpETE (317/215) | 317.2 | 215.1 | 1 | 1 | 10 | 20 | 12 | 15.1 | 30 |
| EPA (301/257) | 301.2 | 257.1 | 1 | 1 | 10 | 16 | 12 | 17.7 | 30 |
| RvD1 (375/141) | 375.2 | 141.1 | 1 | 1 | 10 | 21 | 12 | 9.1 | 30 |

**Supplementary Table 1. Continued**

| **Compound ID** | **Q1** | **Q3** | **Edit Dwell** | **Dwell Time** | **EP** | **CE** | **CXP** | **Expected RT (min)** | **RT Tolerance (s)** |
| --- | --- | --- | --- | --- | --- | --- | --- | --- | --- |
| 17(R)-RvD1 (375/141) | 375.2 | 141.1 | 1 | 1 | 10 | 21 | 12 | 9.3 | 30 |
| RvD2 (375/175) | 375.2 | 175.1 | 1 | 1 | 10 | 30 | 12 | 8.7 | 30 |
| RvD3 (375/147) | 375.2 | 147.1 | 1 | 1 | 10 | 25 | 12 | 8.7 | 30 |
| 17(R)-RvD3 (375/147) | 375.2 | 147.1 | 1 | 1 | 10 | 25 | 12 | 8.5 | 30 |
| RvD4 (375/101) | 375.2 | 101.1 | 1 | 1 | 10 | 22 | 12 | 10 | 30 |
| RvD5 (359/199) | 359.2 | 199.1 | 1 | 1 | 10 | 21 | 12 | 11.9 | 30 |
| PD1 (359/153) | 359.2 | 153.1 | 1 | 1 | 10 | 21 | 12 | 12.1 | 30 |
| PDX (359/153) | 359.2 | 153.1 | 1 | 1 | 10 | 21 | 12 | 11.9 | 30 |
| Maresin 1 (359/221) | 359.2 | 221.1 | 1 | 1 | 10 | 20 | 12 | 12.1 | 30 |
| Maresin 1 (359/250) | 359.2 | 250.1 | 1 | 1 | 10 | 20 | 12 | 12.1 | 30 |
| Maresin 2 (359/221) | 359.2 | 221.1 | 1 | 1 | 10 | 20 | 12 | 13.1 | 30 |
| 4-HDHA (343/101) | 343.2 | 101.1 | 1 | 1 | 10 | 17 | 12 | 16.3 | 30 |
| 7-HDHA (343/141) | 343.2 | 141.1 | 1 | 1 | 10 | 18 | 12 | 15.8 | 30 |
| 13-HDHA (343/193) | 343.2 | 193.1 | 1 | 1 | 10 | 17 | 12 | 15.6 | 30 |
| 14-HDHA (343/205) | 343.2 | 205.1 | 1 | 1 | 10 | 17 | 12 | 15.7 | 30 |
| 16-HDHA (343/233) | 343.2 | 233.1 | 1 | 1 | 10 | 20 | 12 | 15.4 | 30 |
| 17-HDHA (343/245) | 343.2 | 245.1 | 1 | 1 | 10 | 17 | 12 | 15.5 | 30 |
| 7,8-EpDPA (343/113) | 343.2 | 113.1 | 1 | 1 | 10 | 20 | 12 | 16.8 | 30 |
| 10,11-EpDPA (343/153) | 343.2 | 153.2 | 1 | 1 | 10 | 20 | 12 | 16.6 | 30 |
| 13,14-EpDPA (343/161) | 343.2 | 161.1 | 1 | 1 | 10 | 20 | 12 | 16.58 | 30 |
| 16,17-EpDPA (343/274) | 343.2 | 274.1 | 1 | 1 | 10 | 20 | 12 | 16.55 | 30 |
| 19,20-EpDPA (343/241) | 343.2 | 241.1 | 1 | 1 | 10 | 20 | 12 | 16.3 | 30 |
| DHA (327/283) | 327.2 | 283.1 | 1 | 1 | 10 | 18 | 12 | 17.8 | 30 |
| DPA (329/285) | 329.2 | 285.1 | 1 | 1 | 10 | 18 | 12 | 17.9 | 30 |
| Adrenic Acid (331/287) | 331.2 | 287.1 | 1 | 1 | 10 | 18 | 12 | 18 | 30 |
| 9-HODE (295/171) | 295.2 | 171.1 | 1 | 1 | 10 | 25 | 12 | 15 | 30 |
| 13-HODE (295/195) | 295.2 | 195.1 | 1 | 1 | 10 | 25 | 12 | 15.02 | 30 |
| 13-OxoODE (13-KODE) (293/113) | 293.2 | 113.2 | 1 | 1 | 10 | 30 | 12 | 14.84 | 30 |
| 12(13)-EpOME (295/195 | 295.3 | 195.2 | 1 | 1 | 10 | 25 | 12 | 16 | 30 |
| 9,10 DiHOME (313/201) | 313.3 | 201.3 | 1 | 1 | 10 | 30 | 12 | 13 | 30 |
| 10-Nitrolinoleate (324/277) | 324.2 | 277.1 | 1 | 1 | 10 | 18 | 12 | 16.7 | 30 |
| 9(S)HOTrE (293/171) | 293.3 | 171.1 | 1 | 1 | 10 | 25 | 12 | 13.74 | 30 |
| 13(S)HOTrE (293/195) | 293.2 | 195 | 1 | 1 | 10 | 25 | 12 | 13.94 | 30 |
| PAF (508/59) | 508.3 | 59.1 | 1 | 1 | 10 | 20 | 12 | 17.8 | 30 |
| PGE2-d4 (355/193) | 355.2 | 193.1 | 1 | 1 | 10 | 25 | 12 | 8.1 | 30 |
| 15-deoxy-PGJ2-d4 (319/275) | 319.2 | 275.1 | 1 | 1 | 10 | 20 | 12 | 13.6 | 30 |
| 8-iso-PGF2a-d4 (357/197) | 357.2 | 197.1 | 1 | 1 | 10 | 30 | 12 | 7.2 | 30 |
| LTB4-d4 (339/197) | 339.2 | 197.1 | 1 | 1 | 10 | 22 | 12 | 12.1 | 30 |
| LXA4-d5 (356/115) | 356.2 | 115.1 | 1 | 1 | 10 | 19 | 12 | 8.9 | 30 |
| 11,12-EET-d11 (327/171) | 327.2 | 171.1 | 1 | 1 | 10 | 20 | 12 | 16.5 | 30 |
| 5-HETE-d8 (327/116) | 327.2 | 116.1 | 1 | 1 | 10 | 17 | 12 | 15.9 | 30 |
| 12-HETE-d8 (327/184) | 327.2 | 184.1 | 1 | 1 | 10 | 20 | 12 | 15.6 | 30 |
| 15-HETE-d8 (327/226) | 327.2 | 226.1 | 1 | 1 | 10 | 20 | 12 | 15.3 | 30 |
| RvE1-d4 (353/197) | 353.2 | 197.1 | 1 | 1 | 10 | 22 | 12 | 5.9 | 30 |
| RvD2-d5 (380/141) | 380.2 | 141.1 | 1 | 1 | 10 | 23 | 12 | 8.7 | 30 |
| RvD3-d5 (380/152) | 380.2 | 152.1 | 1 | 1 | 10 | 25 | 12 | 8.6 | 30 |
| Maresin 1-d5 (364/221) | 364.2 | 221.1 | 1 | 1 | 10 | 20 | 12 | 12.1 | 30 |
| 6k-PGF1a (369/163) | 369.2 | 163.1 | 1 | 1 | 10 | 35 | 12 | 6.8 | 30 |
| PGE2 and PGD2 (351/189) | 351.2 | 189.1 | 1 | 1 | 10 | 25 | 12 | 8.15 | 30 |
| PGE2 (351/175) | 351.2 | 175.1 | 1 | 1 | 10 | 25 | 12 | 8.06 | 30 |
| 15-keto-PGE2 (349/113) | 349.2 | 113.1 | 1 | 1 | 10 | 28 | 12 | 7.4 | 30 |
| 13,14-dihydro-15-keto PGE2 (351/175) | 351.2 | 175.1 | 1 | 1 | 10 | 26 | 12 | 8.1 | 30 |
| PGD2 (351/233) | 351.2 | 233.1 | 1 | 1 | 10 | 16 | 12 | 8.22 | 30 |
| 11-beta-PGF2a (353/193) | 353.2 | 193.1 | 1 | 1 | 10 | 30 | 12 | 7.5 | 30 |
| 13,14-dihydro-15-keto PGD2 (351/207) | 351.2 | 207.1 | 1 | 1 | 10 | 26 | 12 | 8.8 | 30 |
| PGJ2 (333/271) | 333.2 | 271.1 | 1 | 1 | 10 | 22 | 12 | 10.1 | 30 |
| 15-deoxy-PGJ2 (315/271) | 315.2 | 271.1 | 1 | 1 | 10 | 20 | 12 | 13.6 | 30 |
| PGF2a (353/193) | 353.2 | 193.1 | 1 | 1 | 10 | 34 | 12 | 8.55 | 30 |
| 15-keto PGF2a (351/113) | 351.2 | 113.1 | 1 | 1 | 10 | 35 | 12 | 8 | 30 |
| 13,14-dihydro-15-keto PGF2a (353/291) | 353.2 | 291.1 | 1 | 1 | 10 | 25 | 12 | 9.1 | 30 |
| 20-OH PGF2a (369/193) | 369.2 | 193 | 1 | 1 | 10 | 35 | 12 | 3.22 | 30 |
| TXB2 (369/169) | 369.2 | 169.1 | 1 | 1 | 10 | 22 | 12 | 7.7 | 30 |
| 12-HHT (279/179) | 279.2 | 179.1 | 1 | 1 | 10 | 22 | 12 | 13 | 30 |
| LTB4 (335/195) | 335.2 | 195.1 | 1 | 1 | 10 | 22 | 12 | 12.1 | 30 |
| 20-OH-LTB4 (351/195) | 351.2 | 195.1 | 1 | 1 | 10 | 24 | 12 | 6.6 | 30 |
| 20-COOH-LTB4 (365/195) | 365.1 | 195.1 | 1 | 1 | 10 | 25 | 12 | 6.16 | 30 |
| LXA4 (351/115) | 351.2 | 115.1 | 1 | 1 | 10 | 20 | 12 | 8.9 | 30 |
| 15(R)-LXA4 (351/115) | 351.2 | 115.1 | 1 | 1 | 10 | 20 | 12 | 9.1 | 30 |
| LXB4 (351/221) | 351.2 | 221.1 | 1 | 1 | 10 | 22 | 12 | 8.4 | 30 |
| 5,15-diHETE (335/115) | 335.2 | 115.1 | 1 | 1 | 10 | 22 | 12 | 11.7 | 30 |
| 5S,6R-diHETE (335/115) | 335.2 | 115.1 | 1 | 1 | 10 | 20 | 12 | 13.6 | 30 |
| 5,6 EET (319/191) | 319.2 | 191.1 | 1 | 1 | 10 | 21 | 12 | 16.7 | 30 |
| 8,9 EET (319/155) | 319.2 | 155.1 | 1 | 1 | 10 | 21 | 12 | 16.6 | 30 |
| 11,12 EET (319/167) | 319.2 | 167.1 | 1 | 1 | 10 | 21 | 12 | 16.5 | 30 |
| 14,15 EET (319/219) | 319.2 | 219.1 | 1 | 1 | 10 | 15 | 12 | 16.2 | 30 |

**Supplementary Table 1. Continued**

| **Compound ID** | **Q1** | **Q3** | **Edit Dwell** | **Dwell Time** | **EP** | **CE** | **CXP** | **Expected RT (min)** | **RT Tolerance (s)** |
| --- | --- | --- | --- | --- | --- | --- | --- | --- | --- |
| 5,6 DiHETrE (337/145) | 337.2 | 145.1 | 1 | 1 | 10 | 25 | 12 | 14.4 | 30 |
| 8,9 DiHETrE (337/127) | 337.2 | 127.1 | 1 | 1 | 10 | 30 | 12 | 13.9 | 30 |
| 11,12 DiHETrE (337/167) | 337.2 | 167.1 | 1 | 1 | 10 | 25 | 12 | 13.6 | 30 |
| 14,15 DiHETrE (337/207) | 337.2 | 207.1 | 1 | 1 | 10 | 25 | 12 | 13.4 | 30 |
| 5-HETE (319/115) | 319.2 | 115.1 | 1 | 1 | 10 | 21 | 12 | 15.9 | 30 |
| 9-HETE (319/167) | 319.3 | 167.2 | 1 | 1 | 10 | 21 | 12 | 15.8 | 30 |
| 11-HETE (319/167) | 319.2 | 167.1 | 1 | 1 | 10 | 21 | 12 | 15.4 | 30 |
| 12-HETE (319/179) | 319.2 | 179.1 | 1 | 1 | 10 | 21 | 12 | 15.6 | 30 |
| 15-HETE (319/219) | 319.2 | 219.1 | 1 | 1 | 10 | 19 | 12 | 15.3 | 30 |
| 18-HETE (319/261) | 319.2 | 261.2 | 1 | 1 | 10 | 25 | 12 | 14.74 | 30 |
| 20-HETE (319/289) | 319.2 | 289.1 | 1 | 1 | 10 | 25 | 12 | 14.7 | 30 |
| 8-iso-PGF2a (353/193) | 353.2 | 193.1 | 1 | 1 | 10 | 30 | 12 | 7.2 | 30 |
| AA (303/259) | 303.2 | 259.1 | 1 | 1 | 10 | 16 | 12 | 17.8 | 30 |
| RvE1 (349/195) | 349.2 | 195.1 | 1 | 1 | 10 | 22 | 12 | 5.9 | 30 |
| RvE2 (333/199) | 333.2 | 199.1 | 1 | 1 | 10 | 22 | 12 | 10.1 | 30 |
| RvE2(333/115) | 333.2 | 115.1 | 1 | 1 | 10 | 22 | 12 | 10.1 | 30 |
| 5,15-DiHEPE (333/115) | 333.2 | 115.1 | 1 | 1 | 10 | 22 | 12 | 10.5 | 30 |
| RvE3 (333/201) | 333.2 | 201.1 | 1 | 1 | 10 | 20 | 12 | 12 | 30 |
| LXA5 (349/115) | 349.2 | 115.1 | 1 | 1 | 10 | 20 | 12 | 7.5 | 30 |
| 5-HEPE (317/115) | 317.2 | 115.1 | 1 | 1 | 10 | 19 | 12 | 14.6 | 30 |
| 11-HEPE (317/167) | 317.2 | 167.1 | 1 | 1 | 10 | 19 | 12 | 14.3 | 30 |
| 12-HEPE (317/179) | 317.2 | 179.1 | 1 | 1 | 10 | 19 | 12 | 14.4 | 30 |
| 15-HEPE (317/219) | 317.2 | 219.1 | 1 | 1 | 10 | 18 | 12 | 14.3 | 30 |
| 18-HEPE (317/259) | 317.2 | 259.1 | 1 | 1 | 10 | 16 | 12 | 14.1 | 30 |
| 8,9-EpETE (317/127) | 317.2 | 127.1 | 1 | 1 | 10 | 20 | 12 | 15.4 | 30 |
| 11,12-EpETE (317/167) | 317.2 | 167.1 | 1 | 1 | 10 | 20 | 12 | 15.4 | 30 |
| 14,15-EpETE (317/207) | 317.2 | 207.1 | 1 | 1 | 10 | 20 | 12 | 15.4 | 30 |
| 17,18-EpETE (317/215) | 317.2 | 215.1 | 1 | 1 | 10 | 20 | 12 | 15.1 | 30 |
| EPA (301/257) | 301.2 | 257.1 | 1 | 1 | 10 | 16 | 12 | 17.7 | 30 |
| RvD1 (375/141) | 375.2 | 141.1 | 1 | 1 | 10 | 21 | 12 | 9.1 | 30 |
| 17(R)-RvD1 (375/141) | 375.2 | 141.1 | 1 | 1 | 10 | 21 | 12 | 9.3 | 30 |
| RvD2 (375/175) | 375.2 | 175.1 | 1 | 1 | 10 | 30 | 12 | 8.7 | 30 |
| RvD3 (375/147) | 375.2 | 147.1 | 1 | 1 | 10 | 25 | 12 | 8.7 | 30 |
| 17(R)-RvD3 (375/147) | 375.2 | 147.1 | 1 | 1 | 10 | 25 | 12 | 8.5 | 30 |
| RvD4 (375/101) | 375.2 | 101.1 | 1 | 1 | 10 | 22 | 12 | 10 | 30 |
| RvD5 (359/199) | 359.2 | 199.1 | 1 | 1 | 10 | 21 | 12 | 11.9 | 30 |
| PD1 (359/153) | 359.2 | 153.1 | 1 | 1 | 10 | 21 | 12 | 12.1 | 30 |
| PDX (359/153) | 359.2 | 153.1 | 1 | 1 | 10 | 21 | 12 | 11.9 | 30 |
| Maresin 1 (359/221) | 359.2 | 221.1 | 1 | 1 | 10 | 20 | 12 | 12.1 | 30 |
| Maresin 1 (359/250) | 359.2 | 250.1 | 1 | 1 | 10 | 20 | 12 | 12.1 | 30 |
| Maresin 2 (359/221) | 359.2 | 221.1 | 1 | 1 | 10 | 20 | 12 | 13.1 | 30 |
| 4-HDHA (343/101) | 343.2 | 101.1 | 1 | 1 | 10 | 17 | 12 | 16.3 | 30 |
| 7-HDHA (343/141) | 343.2 | 141.1 | 1 | 1 | 10 | 18 | 12 | 15.8 | 30 |
| 13-HDHA (343/193) | 343.2 | 193.1 | 1 | 1 | 10 | 17 | 12 | 15.6 | 30 |
| 14-HDHA (343/205) | 343.2 | 205.1 | 1 | 1 | 10 | 17 | 12 | 15.7 | 30 |
| 16-HDHA (343/233) | 343.2 | 233.1 | 1 | 1 | 10 | 20 | 12 | 15.4 | 30 |
| 17-HDHA (343/245) | 343.2 | 245.1 | 1 | 1 | 10 | 17 | 12 | 15.5 | 30 |
| 7,8-EpDPA (343/113) | 343.2 | 113.1 | 1 | 1 | 10 | 20 | 12 | 16.8 | 30 |
| 10,11-EpDPA (343/153) | 343.2 | 153.2 | 1 | 1 | 10 | 20 | 12 | 16.6 | 30 |
| 13,14-EpDPA (343/161) | 343.2 | 161.1 | 1 | 1 | 10 | 20 | 12 | 16.58 | 30 |
| 16,17-EpDPA (343/274) | 343.2 | 274.1 | 1 | 1 | 10 | 20 | 12 | 16.55 | 30 |
| 19,20-EpDPA (343/241) | 343.2 | 241.1 | 1 | 1 | 10 | 20 | 12 | 16.3 | 30 |
| DHA (327/283) | 327.2 | 283.1 | 1 | 1 | 10 | 18 | 12 | 17.8 | 30 |
| DPA (329/285) | 329.2 | 285.1 | 1 | 1 | 10 | 18 | 12 | 17.9 | 30 |
| Adrenic Acid (331/287) | 331.2 | 287.1 | 1 | 1 | 10 | 18 | 12 | 18 | 30 |
| 9-HODE (295/171) | 295.2 | 171.1 | 1 | 1 | 10 | 25 | 12 | 15 | 30 |
| 13-HODE (295/195) | 295.2 | 195.1 | 1 | 1 | 10 | 25 | 12 | 15.02 | 30 |
| 13-OxoODE (13-KODE) (293/113) | 293.2 | 113.2 | 1 | 1 | 10 | 30 | 12 | 14.84 | 30 |
| 12(13)-EpOME (295/195 | 295.3 | 195.2 | 1 | 1 | 10 | 25 | 12 | 16 | 30 |
| 9,10 DiHOME (313/201) | 313.3 | 201.3 | 1 | 1 | 10 | 30 | 12 | 13 | 30 |
| 10-Nitrolinoleate (324/277) | 324.2 | 277.1 | 1 | 1 | 10 | 18 | 12 | 16.7 | 30 |
| 9(S)HOTrE (293/171) | 293.3 | 171.1 | 1 | 1 | 10 | 25 | 12 | 13.74 | 30 |
| 13(S)HOTrE (293/195) | 293.2 | 195 | 1 | 1 | 10 | 25 | 12 | 13.94 | 30 |
| PAF (508/59) | 508.3 | 59.1 | 1 | 1 | 10 | 20 | 12 | 17.8 | 30 |

**Supplementary Table 2. Positive Mode**

| **Compound ID** | **Q1** | **Q3** | **Edit Dwell** | **Dwell Time** | **EP** | **CE** | **CXP** | **Expected RT (min)** | **RT Tolerance (s)** |
| --- | --- | --- | --- | --- | --- | --- | --- | --- | --- |

| LTC4-d5 (631/194) | 631.3 | 194.1 | 1 | 1 | 9 | 28 | 15 | 11.5 | 30 |
| --- | --- | --- | --- | --- | --- | --- | --- | --- | --- |
| LTD4-d5 (502/194) | 502.3 | 194.1 | 1 | 1 | 9 | 28 | 13 | 10.2 | 30 |
| LTE4-d5 (445/194) | 445.2 | 194.1 | 1 | 1 | 9 | 23.5 | 13 | 12.32 | 30 |
| LTC4 (626/189) | 626.3 | 189.1 | 1 | 1 | 10 | 28 | 13 | 11.5 | 30 |
| LTD4 (497/189) | 497.3 | 189.1 | 1 | 1 | 10 | 23 | 13 | 10.22 | 30 |
| LTE4 (440/189) | 440.3 | 189.1 | 1 | 1 | 10 | 23 | 13 | 12.3 | 30 |
| MCTR1 (650/191) | 650.3 | 191.1 | 1 | 1 | 10 | 28 | 13 | 11.3 | 30 |
| MCTR2 (521/191) | 521.3 | 191.1 | 1 | 1 | 10 | 23 | 13 | 10.1 | 30 |
| MCTR3 (464/191) | 464.3 | 191.1 | 1 | 1 | 10 | 25 | 13 | 12.24 | 30 |
| PCTR1 (650/231) | 650.3 | 231.1 | 1 | 1 | 10 | 28 | 13 | 11.3 | 30 |
| PCTR2 (521/231) | 521.3 | 231.1 | 1 | 1 | 10 | 23 | 13 | 10.14 | 30 |
| PCTR3 (464/231) | 464.3 | 231.1 | 1 | 1 | 10 | 23 | 13 | 12.3 | 30 |
| PAF (524/184) | 524.3 | 183.9 | 1 | 1 | 10 | 20 | 13 | 17.8 | 30 |
| PGE2 Ethanolamide (PGE2-EA) (396/62) | 396.5 | 62.1 | 1 | 1 | 10 | 25 | 13 | 6 | 30 |
| Oleoyl Ethanolamide (OEA) (326/62) | 326.4 | 62.1 | 1 | 1 | 10 | 25 | 13 | 17.9 | 30 |
| Palmitoyl Ethanolamide (300/62) | 300.4 | 62.1 | 1 | 1 | 10 | 25 | 13 | 16.6 | 30 |
| Anandamide (AEA) (348/62) | 348.4 | 62.1 | 1 | 1 | 10 | 25 | 13 | 17.6 | 30 |
| Docosahexaenoyl Ethanolamide (DHEA) (372/62) | 372.4 | 62.1 | 1 | 1 | 10 | 25 | 13 | 17.7 | 30 |
| Linoleoyl Ethanolamide (LEA) (324/62) | 324.4 | 62.1 | 1 | 1 | 10 | 25 | 13 | 17.5 | 30 |
| Stearoyl Ethanolamide (ceramid) (328/62) | 328.4 | 62.1 | 1 | 1 | 10 | 25 | 13 | 18 | 30 |
| oxy-Arachidonoyl Ethanolamide (oxy-AEA) (364/62) | 364 | 62 | 1 | 1 | 10 | 25 | 13 | 14.5 | 30 |
| 2-Arachidonoyl Glycerol (2AG) (379/287) | 379.4 | 287.2 | 1 | 1 | 10 | 25 | 13 | 17.73 | 30 |
| Docosatetraenoyl Ethanolamide (DEA) (376/62) | 376.6 | 62.1 | 1 | 1 | 10 | 25 | 13 | 17.9 | 30 |
| alpha-linolenoyl ethanolamide (322/62) | 322.4 | 62.1 | 1 | 1 | 10 | 25 | 13 | 16.4 | 30 |
| oleamide (282/247) | 282.4 | 247.4 | 1 | 1 | 10 | 25 | 13 | 17.9 | 30 |
| dihomo-gamma-linolenoyl ethanolamide (350/62) | 350.4 | 62.1 | 1 | 1 | 10 | 25 | 13 | 17.8 | 30 |
| docosanoyl ethanolamide (384/62) | 384.5 | 62.1 | 1 | 1 | 10 | 25 | 13 | 19 | 30 |

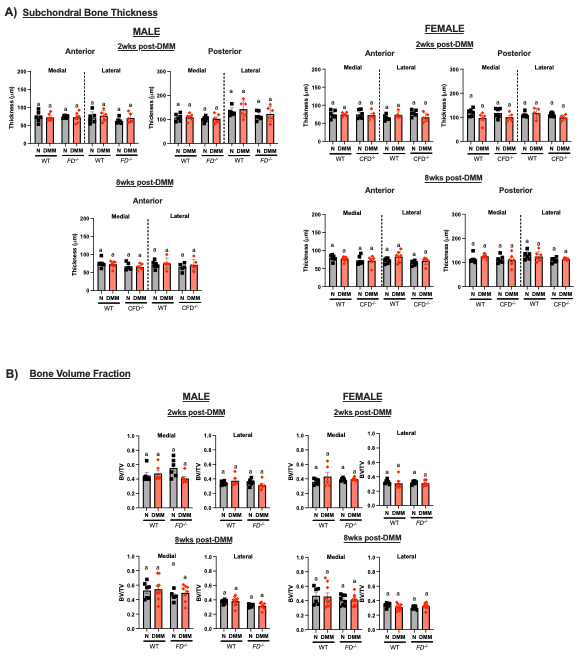

**Supplementary Figure 1. There were no other changes to bone.** (A) There were no differences in subchondral bone thickness at 2 weeks post-DMM in male mice. There were no differences in female due to surgery or strain. B) Bone volume fraction (BV/TV) was not significantly different between strain, surgery, or sex in the proximal tibial epiphysis.

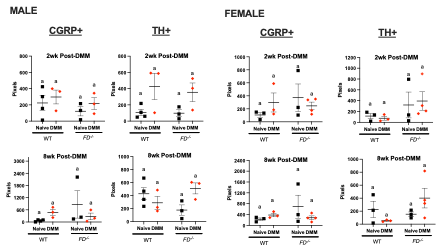

**Supplementary Figure 2. There were no differences in number of CGRP+ and TH+ nerve endings.** Preliminary analysis of (a) CGRP+ and (b) TH+ neurites showed no differences in between strain, sex, or surgery at the joint.

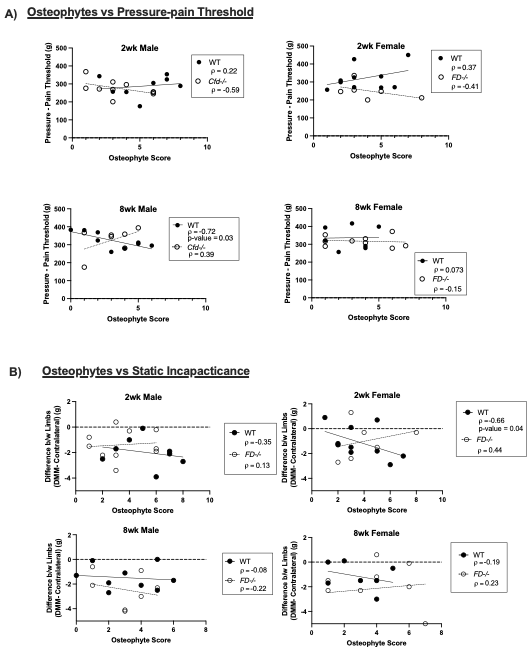

**Supplementary Figure 3. There are no correlations between osteophytes and pressure-pain threshold or static incapacitance.** (A) Spearman’s correlation between osteophytes and pressure-pain threshold. (B) Spearman’s correlation between osteophytes and offloading of the surgical limb. Spearman’s correlation coefficient (ρ) is reported. p-value > 0.05 is indicated in the graphs.

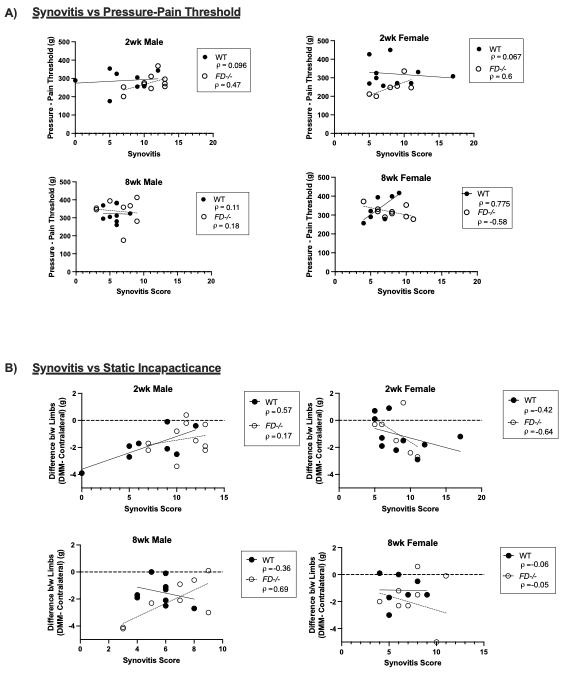

**Supplementary Figure 4. There are no correlations between synovitis and pressure-pain threshold or static incapacitance.** (A) Spearman’s correlation between synovitis and pressure-pain threshold. (B) Spearman’s correlation between synovitis and offloading of the surgical limb. Spearman’s correlation coefficient (ρ) is reported. p-value > 0.05 is indicated in the graphs.

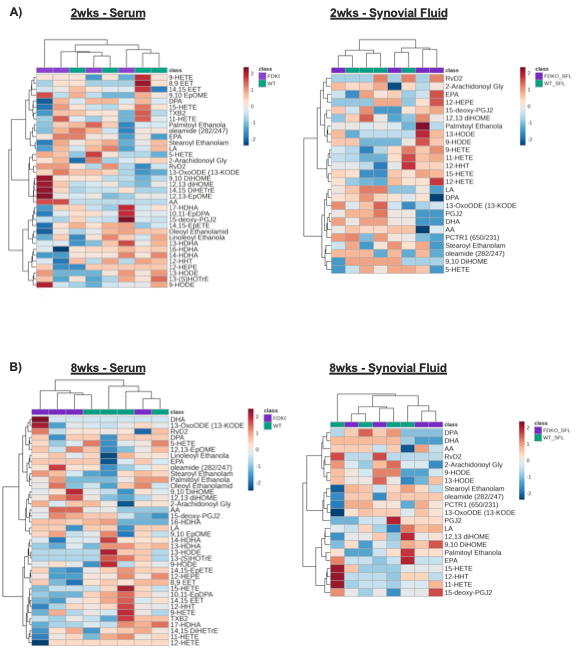

**Supplementary Figure 5. Heat maps of targeted lipidomic profiles of *FD^-/-^* and WT DMM groups.** (A) 2 weeks and (B) 8 weeks post-DMM in male mice.

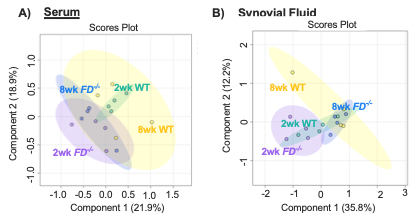
**Supplementary Figure 6. PLS-DA plot comparing strain and timepoint separated into four groups.** (A) Serum (B) Synovial fluid of male mice.

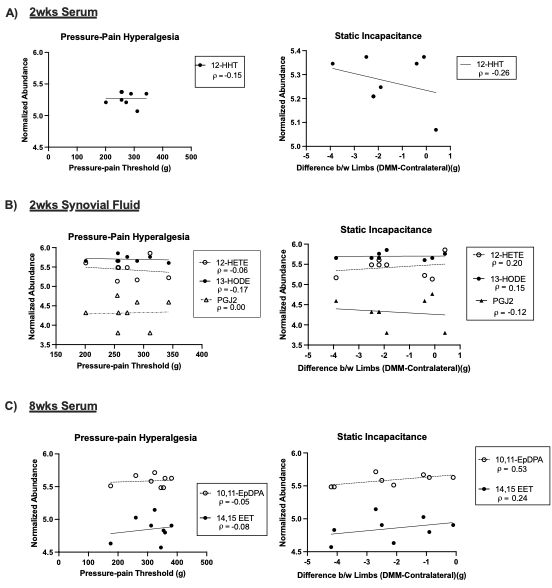

**Supplementary Figure 7. There are no correlations between pressure-pain threshold and abundances of significant eicosanoids.** (A) Serum at two weeks (B) Synovial fluid at two weeks (C) Serum at 8 weeks post-DMM. Spearman’s correlation coefficient (ρ) is reported.
